# LNCRNA expression landscape and specificity between brain regions

**DOI:** 10.1101/2021.10.29.466410

**Authors:** Ogunleye Adewale Joseph, Umair Ali, Olufemi Michael Juwon

## Abstract

Long noncoding RNAs (lncRNAs) are transcribed into low potential protein coding RNA molecules, which account for over 70% of mammalian transcriptional products. The role of lncRNAs and their expression is still largely unknown, and the subject of recent investigations. Here, we used bulk RNA sequencing data from the Genotype-Tissue Expression (GTEx) project to reveal the occurrence and identify the specificity of lncRNAs in 13 brain regions (1000 samples). We observed that these highly specific lncRNA were co-expressed with previously known mRNA markers for the 13 study regions of the brain. Further investigation revealed that splicing could influence the divergent biogenesis and enrichment of specific lncRNA alleles in different brain regions. Overall, we demonstrate the use of lncRNA as an independent tool for deconvolving brain regions and further highlights its use for cell-type identification from bulk transcriptome data.

## Introduction

LncRNA are transcriptional products with >200 nucleotides without a distinct open reading frame and poor protein coding potential [1,2]. LncRNA can regulate major cellular processes such as transcription and epigenetics by initiating signal transduction [3,4], sponging microRNA [5–7], acting as a scaffold for assembly molecules and guiding proteins to their targets [1,8,9]. Unlike messenger RNA (mRNA), lncRNA are localized in the nucleus where they directly interact with the genome to influence overall cell development [2,10]. It is believed that lncRNAs are involved in modulating gene expression,and their canonical functions start at a genetic locus. However, lncRNAs are transcribed from enhancers,called eRNAs.both are heterogenous molecules and different from each other but perform the same function in the progress of mRNA transcription and translation [11–14]. However lncRNAs do not possess protein coding ability, spatiotemporal-specific expression patterns have enhanced the different functions and complex processes of lncRNAs [15]. For the functionality of lncRNAs the precise processing, tissue specificity, subcellular localization and differential expression of lncRNAs can be used as arguments. It could be taken logically as that these observations are also consistent with lncRNAs being the product of noisy transcription. LncRNAs exhibit tissue-type, cell-type and cell-type condition specificity in their expression pattern. Generally, about 11% of lncRNAs are ubiquitous in all human tissues [16]. LncRNAs are abundantly expressed in the brain, however, the tissues have been shown to be non-overlapping in their expression patterns [17,18]. For example, high throughput RNA in-situ hybridization analysis of the Allen Brain Atlas (adult mouse brain) [19,20] identified 849 out of 1328 lncRNAs to be associated with a distinct brain region, cell or subcellular compartment [21]. Due to the apparent profile of distinct lncRNA expression, brain regions such as hippocampus, cerebral cortex, olfactory bulb and cerebellum, some studies have proposed lncRNAs as biomarkers for some brain regions [17,18,22]. It is also safe to assume that this distinct specificity within regions suggests that lncRNA may play a role in the development and homeostasis of brain regions as opposed to the popular notion that they are transcriptional noise. Resolving the population and abundance of lncRNA within brain regions poses several fundamental constraints. The poor inter-species conservation of lncRNA raises evolutionary appreciation but may contradict extrapolation of functional insights. Furthermore, lncRNAs show transient expression between developmental stages [18,23] which may nullify the significance of a comprehensive catalogue. In this study, we attempted to resolve the distinct population of lncRNAs between 13 specific regions in 1000 adult human brains RNA samples as documented by the GTEx consortium [22,24,25]. The enrichment of different lncRNAs within variable brain regions was apparent. Since the GTEx data also provides a significant resource on splicing, we investigated possible contributions of the special splicing program of the brain to lncRNA expression.

## Methods

### Bulk RNA-Seq data preprocessing

Bulk RNA-Seq data (measured in transcript per million; TPM) was retrieved from the Genotype Tissue Expression (GTEx) consortium (https://gtexportal.org/home/datasets) [22,24,25]. For the purpose of this study, we randomly selected 1000 brain samples containing 13 brain regions (cortex, frontal cortex, nucleus accumbens, caudate, cerebellar hemisphere, cerebellum, putamen, hypothalamus, substantia nigra, hippocampus, spinal cord, anterior singlet cortex and amygdala) as classified GTEx. 5885 lncRNAs were selected based on the annotations in GENCODE release 32 [26,27]. Due to the unstable nature of lncRNA nomenclature, we decided to stick with the much more ensembl ID.

### Cluster analysis

We performed semi-supervised clustering to group lncRNAs into distinct expression clusters using Uniform Manifold Approximation and Projection and manual cluster labelling. First, we removed lncRNAs with non-variable expressions, and performed tissue-wise count normalization. We implemented UMAP within ScanPy [28] for 1754 lncRNAs and recovered 13 biologically relevant clusters with non-repetitive transcript profiles. Cluster analysis presents a unique challenge associated with defining the biological identity of each cluster. This was solved by ranking lncRNAs within each cluster using Wilcoxon’s rank sum analysis after which we assigned labels manually using GTEx transcript browser.

### Splicing across regions

Generally, lncRNAs are weakly spliced, but such rare events are known to contribute to vary within tissues [29,30]. Splicing quantitative trait loci (sQTL) (minor allele frequency) data was retrieved from GTEx and preprocessed for the lncRNAs expressed in the 13 brain regions. To estimate the variation of splicing between regions, we performed two independent analyses; (i) We inferred lncRNA trajectories between the regions and the trajectory pseudotime using partition-based graph abstraction (PAGA). (ii) We performed hierarchical clustering for each region based on the presence and abundance of lncRNA splicing alleles and ranked the outputs. The ranked output of each of these independent analyses was manually compared.

### Cerebellar cell-type identification

Upon identification of the highly variable lncRNAs in each region, we performed pairwise correlation analysis to determine co-expression pattern between these lncRNAs and 4 highly expressed mRNAs markers in each of the 4 cerebellar cells (Granule-cell, Purkinje-cell, Astrocyte, Oligodendrocyte) identified by colorimetric in-situ hybridization experiments from the Allen brain atlas (ABA) [20,31]. ABA provides at least 5 cell-specific mRNA markers for each cell; granule cell (*Zranb2, Capn7, Mfap3, Ubr1* and *Tiam1*), purkinje cell (*Car8, Large, Pvalb, Gsbs* and *Lmo7*), astrocyte (*Dao1, Tmem47, Mlc1, Sox21* and *120009O22Rik*) and oligodendrocyte (*Hspa2, Anln, Mobp, Mylk* and *Klk6*) [31]. We collected count data for all 20 mRNA from the same GTEx dataset we are using. Upon pairwise correlation, we used k-mean clustering to iteratively identify the maximum number of biologically relevant clusters that can be extracted from the correlation matrix. Pairwise correlation was performed with pandas corr function, k-mean clustering and visualization was done in ggplot2 (R).

## Results and Discussion

### There is a gradual transition in the population lncRNAs in anatomically close brain regions

Characterizing tissue-specific expression of lncRNA is critical to understanding their phenotypic consequences. In the current study, we deconvolved 13 brain regions using bulk lncRNA expression data from 1000 de-identified brain regions to understand the relationship between lncRNA and observed phenotypes. Upon data preprocessing, we focused on 1750 out of 5885 lncRNAs with hypervariable expression profiles to describe region-specific transcription profiles. Using Uniform Manifold Approximation and Projection (UMAP) embedding, we identified 13 clusters based on lncRNA expression count for 1000 brain regions (Figure 2a). To confirm that these clusters are biological correspondents of the brain regions, we ranked all lncRNAs in the cluster using p-value estimates from Wilcoxon sum rank analysis (<u>Supplementary Table 1</u>). We then manually queried the GTEx transcript browser using the top 5 ranked lncRNA in each cluster to reflect the true biological identity.

**Figure 1:**
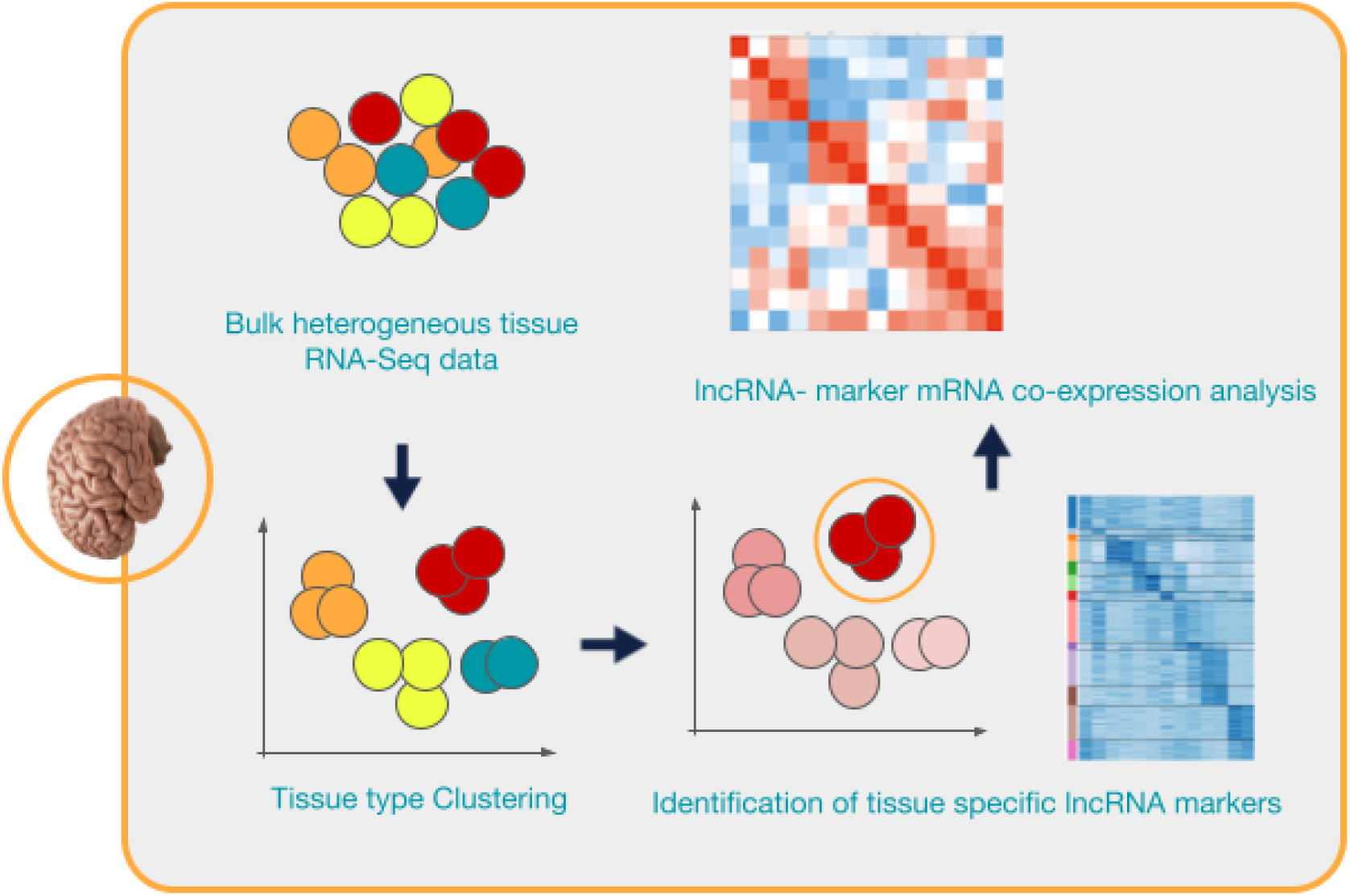
Workflow for the methods used in this study. We collected data from the GTEx dataset, performed UMAP clustering, manually identified the biological identity of the clusters and finally, used co-expression correlation to identify four cell types in the cerebellum.

**Figure 2:**
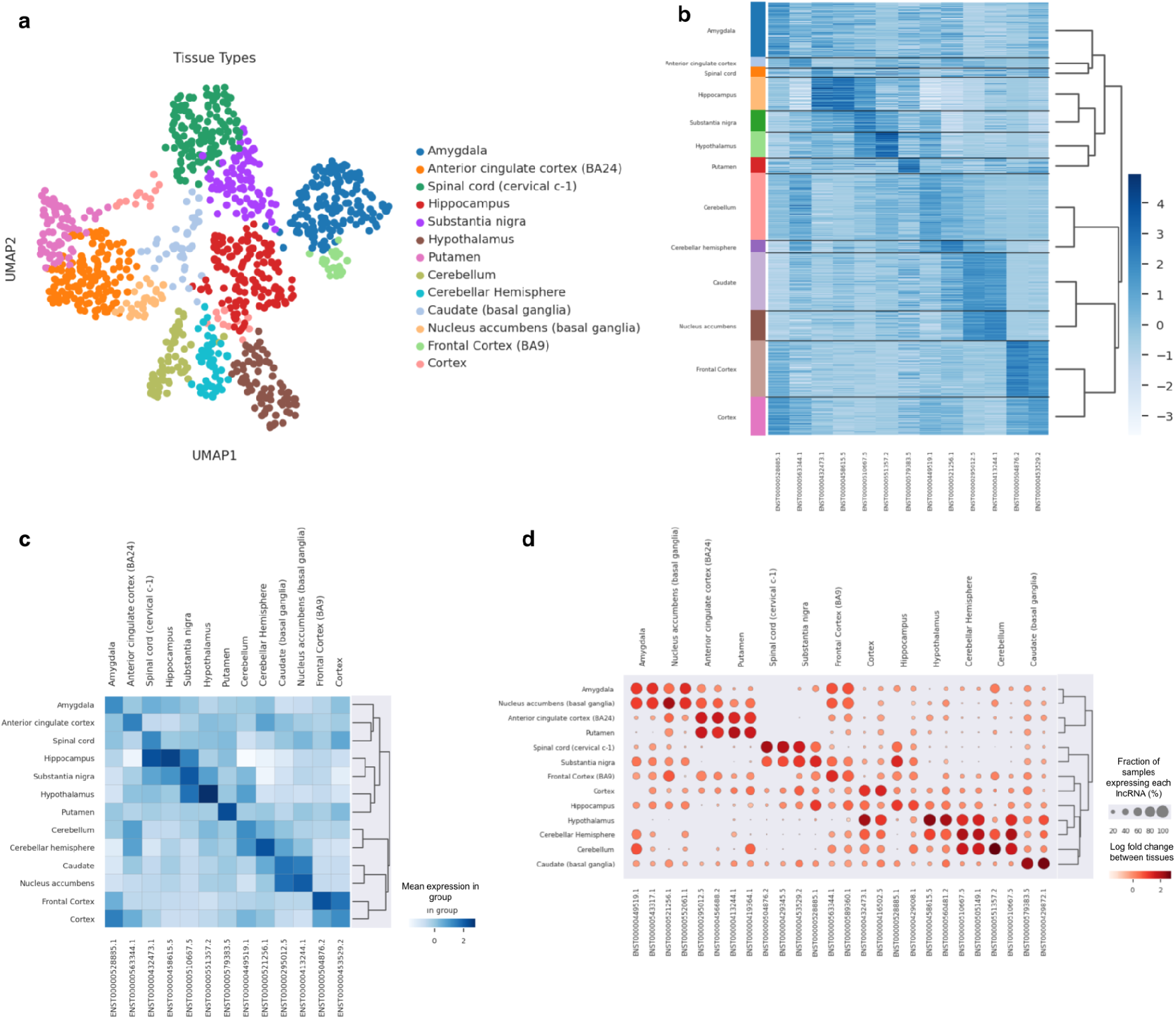
(a) Clustering and identification of the 13 brain regions from UMAP calculations from 1000 de-identified samples and 1750 lncRNAs. (b) Heatmap showing the enrichment in expression of each brain region marker based on Log-transformed normalized expression. (c) Mean expression of each lncRNA marker compared over all the brain region clusters. (d) Log2 fold change of the top 2 lncRNA markers between the different brain regions and the percentage of the number of samples expressing each lncRNA marker in each group. (NB: Due to the unstable nature of lncRNA nomenclature, we decided to stick with the much more ensembl ID).

We were able to detect a small population of lncRNA (usually 3-5) that are really specific to each region. We further observed a gradual but discrete shift in the presence and abundance of lncRNAs between the different brain regions. This is particularly obvious for regions that are anatomically close such as cerebellum and cerebellar hemisphere, cortex and frontal cortex and brain regions in the basal ganglia (substantia nigra, hypothalamus and putamen). This might suggest a spatial link in the abundance and gradual spread of lncRNA over closely associated regions of the brain (Figure 2b-d). In a recent preprint performed by [32], an unsupervised hierarchical clustering to segregate the brain regions based on the transcriptome of the chimpanzee’s brain. In an attempt to correlate the transcriptome transitions with evolutionary development of brain regions, they recovered putamen, substantia nigra and hypothalamus in this order. The ability of lncRNA to also provide a similar transcriptome transition between the regions suggests a role for lncRNA in their differentiation.

Also, some ubiquitously expressed lncRNAs such as *MEG3, OBSCN, ZFAS1, SNHG15* and *PART1* were found as part of the top ranked genes in many clusters. Previous studies have shown that ubiquitous lncRNAs are important for basal cell function and some house-keeping roles [33]. We suggest that they may cooperate with the singly expressed lncRNAs to stabilize the core lncRNA network that defines each brain region.

### Splicing

LncRNAs are known to be rarely spliced due to their weak internal splicing signals which is a consequence of the relatively longer distance between the 5’ splice site, 3’ splice sites and branch point [22,29]. Nevertheless, a few lncRNA splicing variants have been identified and they correlate with tissue variation. Indeed, we retrieved 109 lncRNAs with splice allele variants (minor alleles) across all brain regions from the GTEx database which agrees with the low splicing frequency of lncRNAs. We created a minor allele frequency (MAF) matrix between these 109 lncRNA and the 13 brain regions. To capture the influence of lncRNA splicing, 2 independent strategies were performed using the MAF matrix. First, we selected the cerebellum as an initial root to infer a developmental trajectory and prediction of the pseudo-temporal transition lncRNAs between the brain regions based on the 1750 hypervariable lncRNAs. Secondly, we performed ranked hierarchical clustering of the MAF matrix (109 lncRNAs). We then manually compared the two ranked outputs in a pairwise matrix. Although we could not infer a direct cause-effect relationship between splicing abundance and tissue variation, our analysis showed that brain regions with low splicing allele frequencies (substantia nigra, frontal cortex, amygdala, spinal cord and putamen) were terminal endpoints of the trajectory analysis and hierarchical tree (figure 3a-c). Implicitly, it can be assumed that the trajectories represent the differentiation of brain regions on a time scale, as well as a qualitative and quantitative transition in the estimate of lncRNAs splicing across different brain regions.

**Figure 3:**
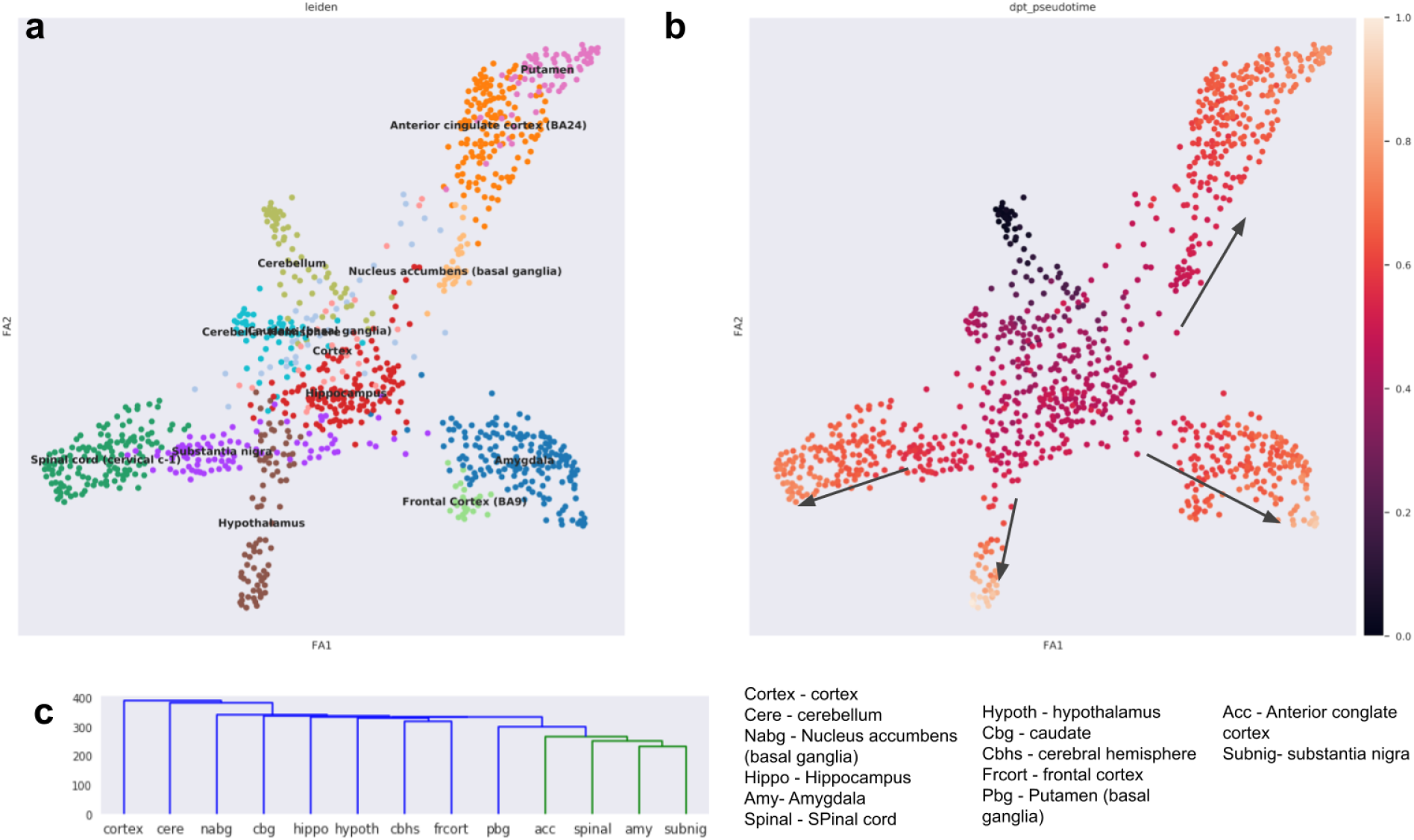
(a) trajectory inference reflects the pseudo-temporal transition and connectivity of lncRNAs between brain regions (b) Trajectory pseudotime analysis (with cerebellum as a starting point) shows that substantia nigra, frontal cortex, amygdala, spinal cord and putamen as endpoints of the trajectory. (c) Anterior cingulate cortex, substantia nigra, amygdala and spinal cord are terminal endpoints of the hierarchical clustering. Compared with the observation of the trajectory analysis, they are terminal endpoints of the pseudotime prediction.

### Cerebellar cell-type identification via mRNA-lncRNA co-expression analysis

The cerebellum regions were richly represented in the GTEx dataset, with distinct lncRNA markers. Using pairwise correlation analysis, we identified co-expression patterns between mRNA markers (retrieved from ABA, see methods) and total lncRNA transcriptome of the cerebellum. We further dissected the correlation matrix into 4 clusters (k-mer, see methods) that matched each cell type (figure 4). The identity of the clusters were verified by the mRNA marker in each cluster. Overall, we identified lncRNA markers for granule cells (lncRNA: *RP11-350N15*.*4-001, RP3-405J10*.*3-001*), purkinje cells (lncRNA: *LINC00304-001, LINC00304-002, RP1-310O13*.*7-001, RP13-895J2*.*2, RP13-895J2*.*2-002*), astrocytes (lncRNA: *AC005786*.*7, RP11-23P13*.*6-001, RP11-23P13*.*6-002*) and oligodendrocytes (lncRNA: *MEG3-026, MEG3-004*,) (see <u>supplementary table 2</u>).

**Figure 4:**
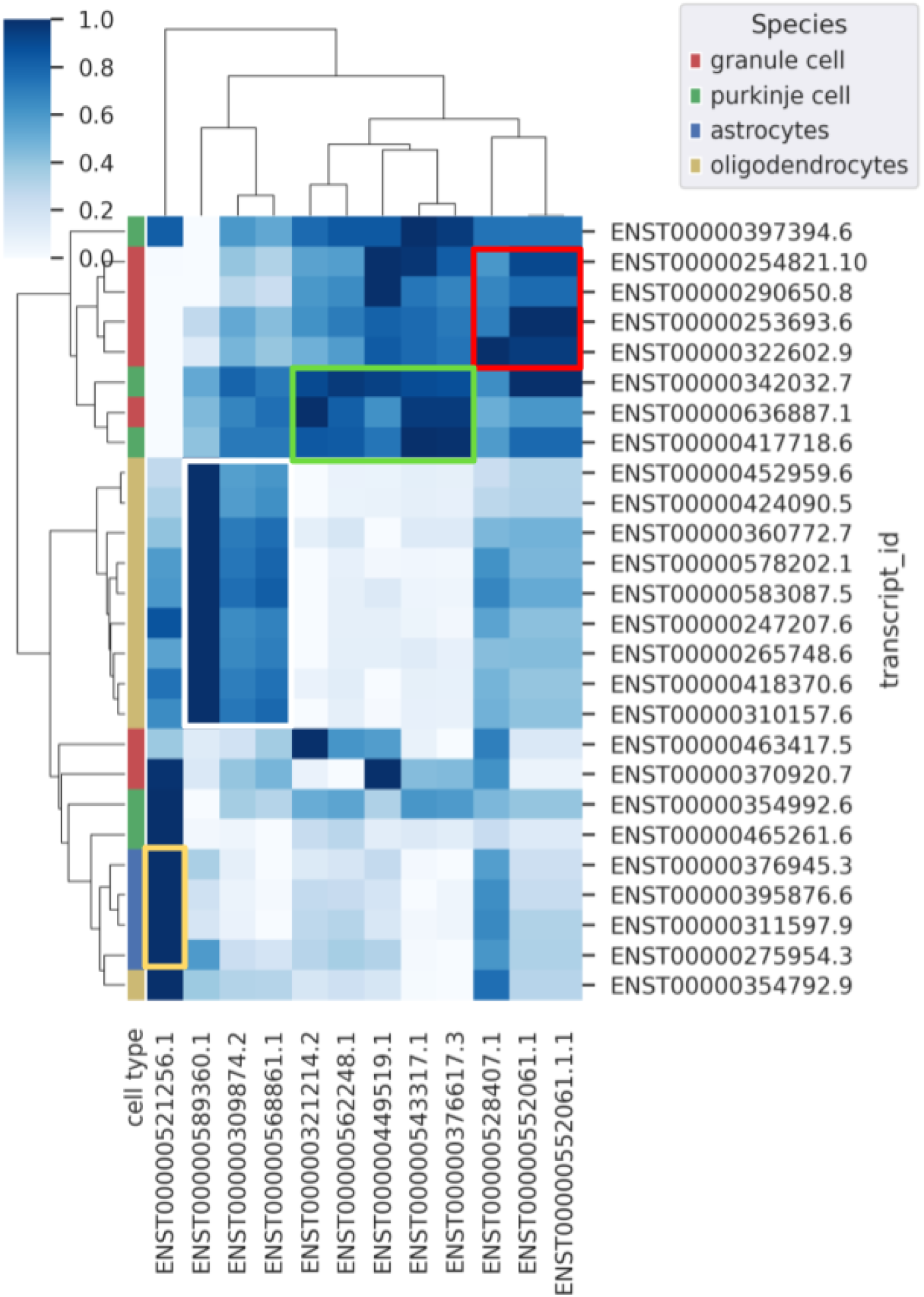
Co-expression analysis of the lncRNA and marker mRNAs predicts four cerebellar cell-types; granule cells (red box), purkinje cells (green box), astrocytes (white box) and oligodendrocytes (yellow box).

Of important note is the expression and specificity of MEG3 and its variants to cerebellar oligodendrocytes. Previous studies have noted the absence of MEG3 in the nucleus of mature oligodendrocytes. However, a recent study performed by [34], identified 3 distinct populations of oligodendrocyte progenitor cells (OPCs; OPC1, OPC2 and OPC3) with MEG3 expression in only OPC3. Our results provide supporting evidence and suggest that MEG3 expression may be crucial to OPC lineage migration into the cerebellum. The functional significance of *LINC00304* and its variants is currently not well understood. Nevertheless, we detected its high expression in the cerebellar cortex of the developing human brain atlas as curated by the harmonizome database [35], while [36] profiled it as a regulator of CpG specific DNA methylation in obesity patients. Unfortunately we could not profile specific roles for astrocyte and granule cell-specific lncRNAs. Overall, we reflect the essentiality of lncRNAs as a sufficient molecular tool for deconvolving the cell types even from bulk lncRNA data.

## Conclusion

Overall, we investigated the expression of lncRNA with respect to their regional specificity in the brain. We showed that some subset of lncRNA could efficiently form a network that can define the identity of each brain region. We further leveraged on the cell specificity of lncRNAs to predict four cell types in the cerebellum. We propose that future studies should verify the functional modularity of these lncRNAs to cell physiology using targeted approaches.

## Abbreviations

lncRNA: Long non coding Ribonucleic acid
RNA: Ribonucleic acid
DNA: Deoxyribonucliec acid
mRNA: Messenger Ribonucleic acid
GTEx: Genotype-Tissue Expression
MAF: Minor allele frequency
sQTL: Splicing quantitative trait loci
PAGA: Partition-based graph abstraction

## Conflicts of interest

None to be declared.

## Notes

### Competing Interest Statement

The authors have declared no competing interest.

